# Heterochromatin Interactions Maintain Homologous Centromere Associations in Mouse Spermatocyte Meiosis

**DOI:** 10.1101/264432

**Authors:** Hoa H. Chuong, Craig Eyster, Chih-Ying Lee, Roberto J. Pezza, Dean Dawson

## Abstract

In meiosis, crossovers between homologous chromosomes link them together. This enables them to attach to microtubules of the meiotic spindle as a unit, such that the homologs will be pulled away from one another at anaphase I. Homologous pairs can sometimes fail to become linked by crossovers. In some organisms, these non-exchange partners are still able segregate properly. In several organisms, associations between the centromeres of non-exchange partners occur in meiotic prophase. These associations have been proposed to promote segregation in meiosis I. But how centromere pairing could promote subsequent proper segregation is unclear. Here we report that meiotic centromere pairing if chromosomes in mouse spermatocytes allows the formation of an association between chromosome pairs. We find that peri-centromeric heterochromatin connections tether the centromeres of chromosome pairs after dissolution of centromere paring. Our results suggest that, in mouse spermatocytes, heterochromatin maintains the association of chromosome centromeres in the absence crossing-over.

## Introduction

Faithful homologous chromosome segregation at the first meiotic division depends upon connections that tether homologous chromosome pairs. The connections are normally created by crossovers between the homologous partners (Bascom-Slack et al. 1997). Chiasmata, the cytological manifestation of crossovers, keep the partners connected as they become stably oriented on the metaphase spindle. Stable attachments are formed when opposing microtubules pull the partner chromosomes towards opposite poles of the spindle creating tension that stabilizes the kinetochore-microtubule attachments (Nicklas and Koch 1969; Nicklas 1997). This tension is transmitted across the connection created by the chiasma nearest to the centromeres. Consequently, mutations that eliminate recombination are invariably associated with increased errors during meiotic chromosome segregation (Klapholz et al. 1985; Dernburg et al. 1998; Klein et al. 1999; Baudat et al. 2000; Romanienko and Camerini-Otero 2000), reviewed in (Sansam and Pezza 2015). However, individual chromosome pairs that have failed to become joined by crossovers can nonetheless segregate properly in some organisms. In *Drosophila* and yeast a high proportion of non-exchange chromosomes (those without crossovers) partition correctly in meiosis I (GRELL 1962; Dawson et al. 1986; Hawley et al. 1992; Davis and Smith 2003). Thus, these organisms have mechanisms, beyond crossing-over, that can promote proper meiotic disjunction. There are suggestions that this may also be the case in mammals. In mice, the majority of chromosomes in oocytes from a recombination-deficient mutant appeared to be spatially balanced on the spindle, as if there are mechanisms to partition equal numbers of chromosomes to each pole (albeit not the correct chromosomes) (Woods et al. 1999). In humans, while smaller chromosomes (21 and 22) fail to experience crossovers in about 5% of meioses (Oliver et al. 2008; Fledel-Alon et al. 2009; Cheng et al. 2009), they are estimated to non-disjoin in only <1% of meioses (Tease et al. 2002; Oliver et al. 2008; Fledel-Alon et al. 2009). Therefore, it maybe that non-disjunction in mammals, as in yeast and Drosophila, may reflect the failure of multiple mechanisms: first, failure to generate a crossover, and second, failure of one or more back-up systems that promote proper segregation of achiasmate (non-exchange) partners (Oliver et al. 2008; Fledel-Alon et al. 2009; Cheng et al. 2009).

Mechanisms that partition non-exchange chromosome partners have been described in yeast and *Drosophila*. In these, organisms the centromeres of non-exchange chromosomes pair or cluster in meiotic prophase (Ding et al. 2004; Gladstone et al. 2009; Newnham et al. 2010; Takeo et al. 2011). In budding yeast, centromere pairing in meiotic prophase predisposes the non-exchange partners to segregate properly in anaphase and may also contribute significantly to the segregation fidelity of recombined chromosomes (Kemp et al. 2004; Gladstone et al. 2009; Newnham et al. 2010).

The manner by which prophase centromere pairing in these organisms promotes disjunction is unclear. Homologous partners become tightly aligned in meiotic prophase by a structure called the synaptonemal complex (SC) that runs along their aligned axes. In budding yeast, the SC disassembles from the chromosome arms in late prophase except at the centromeres where it mediates their pairing (Gladstone et al. 2009; Newnham et al. 2010). But this centromeric SC largely disappears before metaphase when chromosomes become attached to the microtubules (Gladstone et al. 2009), so the pairing it provides cannot be the basis for mediating bi-orientation of the centromeres on the spindle. In *Drosophila*, segregation of non-exchange partners also appears to depend on pairing of their centromeric regions in prophase (Karpen et al. 1996; Dernburg et al. 1996). Observations of non-exchange chromosome partners in metaphase I in *Drosophila* oocytes show that the centromeres are not directly paired during the bi-orientation process, but instead may be connected by threads of pericentromeric heterochromatin (Hughes et al. 2009). Together, these results suggest the model that tight centromere pairing in prophase may allow the formation of chromatin connections that can then promote bi-orientation in metaphase.

In mouse spermatocytes, homologous partners experience a period of prophase centromere pairing (Bisig et al. 2012; Qiao et al. 2012). As in budding yeast, the pairing is mediated by SC components at the centromeres after SC disassembly, and the pairing dissolves before prometaphase. As in most eukaryotes, the centromeres of mouse chromosomes are flanked by blocks peri-centromeric heterochromatin (Pardue and Gall 1970; Mouse Genome Sequencing Consortium et al. 2002; Martens et al. 2005). In early meiotic prophase, the pericentromeric regions of chromosomes associate in clusters called chromocenters (Schwarzacher et al. 1984; Scherthan et al. 1996; Berríos et al. 2010; Takada et al. 2011; Gómez et al. 2013). Multiple centromeres cluster in each chromocenter (Berríos et al. 2010; Berríos et al. 2014; Hopkins et al. 2014), with homologous centromeres usually in different chromocenters (Takada et al. 2011). Thus, the mechanism that clusters the centromeres is homology-independent.

Here we demonstrate that synaptonemal complex formation, re-orders pericentromeric associations, helping homologous centromeres move to the same chromocenters where they become tightly paired by SC components. After the SC-mediated centromere pairing dissolves in late prophase, the pericentromeric heterochromatin masses of the homologous partners remain associated, keeping homologous centromeres linked, even for chromosomes apparently not tethered by chiasmata. Together these observations suggest a mechanism by which centromere pairing in prophase might promote the segregation of non-exchange partners at anaphase I.

## Results

### Pericentromeric heterochromatin moves from non-homologous to homologous associations through meiotic prophase

We monitored the behavior of peri-centromeric heterochromatin in mouse spermatocytes to explore the possibility that interactions of the heterochromatin regions might promote proper meiotic chromosome segregation. All mouse chromosomes are sub-telocentric (the centromere is near one telomere), with a block of peri-centromeric heterochromatin adjacent to the end that harbors the centromere (Pardue and Gall 1970; Mouse Genome Sequencing Consortium et al. 2002; Martens et al. 2005). Mice have 20 pairs of chromosomes (19 pairs of somatic chromosomes and an XY pair in males), thus complete pairing of homologous centromeres in pachytene spermatocytes would yield twenty-one centromeric signals (nineteen autosome pairs plus the X and Y), while completely dispersed centromeres would yield forty signals. We scored the number of heterochromatin signals (DAPI) in wild-type cells at advancing meiotic stages (S-phase through late prophase) (Fig. 1 A). Centromeres were identified by their characteristic knob of SYCP3 staining (Moens and Spyropoulos 1995; Parra 2004) and by staining with CREST antibodies that recognize centromere proteins. As described previously (Berríos et al. 2010; Hopkins et al. 2014), from pre-meiotic stages through prophase the centromeres cluster in chromocenters (Fig. 1 A and Fig. S1). In early prophase (leptotene) there are fewer and larger chromocenters containing high numbers of centromeres. By diplotene the chromocenters have resolved to become smaller and significantly more numerous, harboring fewer centromere pairs ((Fig. 1 A, Fig. S1). In mid-diplotene, centromeres are usually tightly paired with their partners by a short remnant of persisting synaptonemal complex (Fig. 1 A, diplotene, white arrowheads) (Bisig et al. 2012; Qiao et al. 2012). By late diplotene this tight centromere paring dissolves, but the homologous centromere pairs remain within the same chromocenter, sometimes together with the centromeres of other homologous pairs (Fig. 1 B).

**Figure 1.**
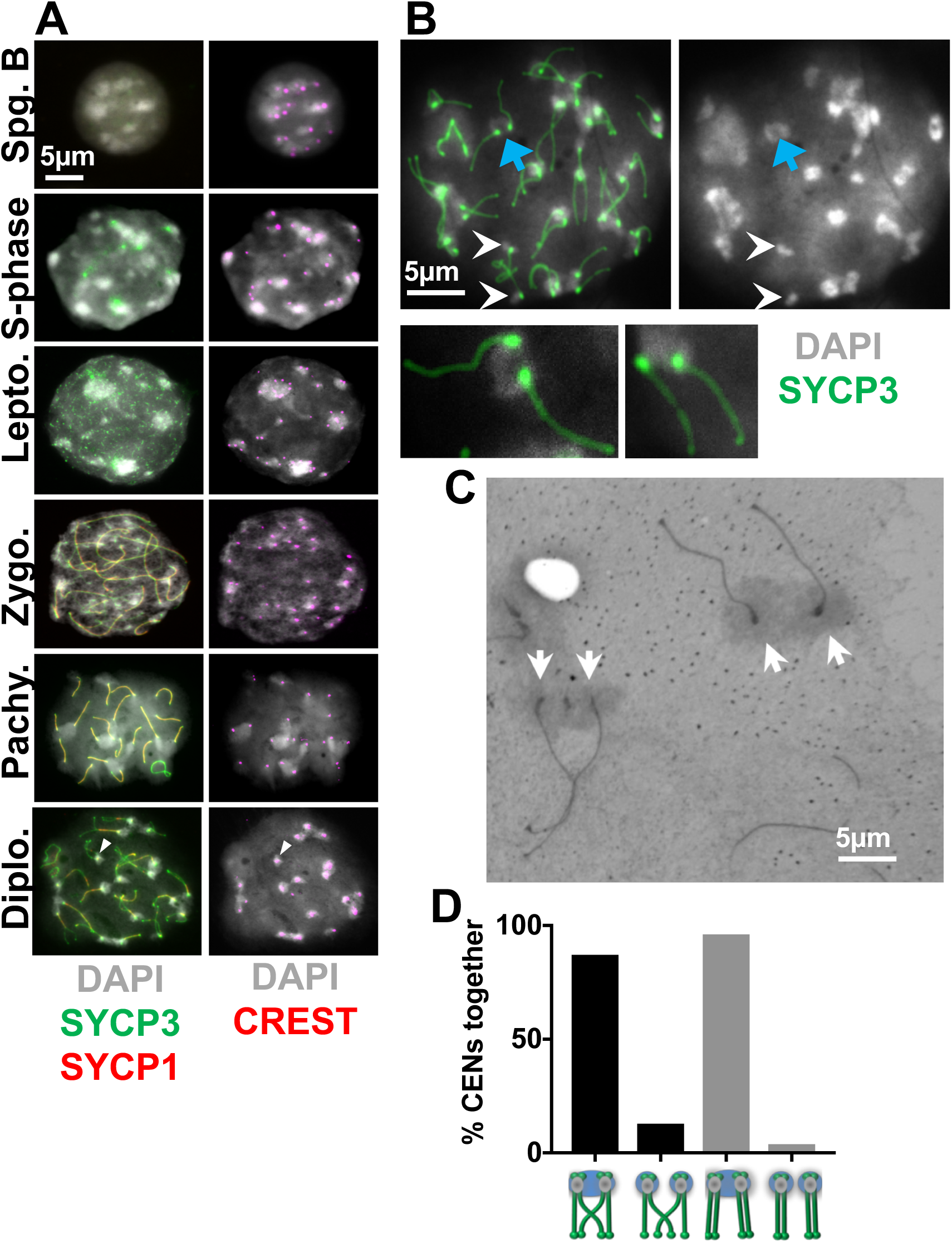
Dynamics of centromeric heterochromatin configuration during prophase in mouse spermatocytes. **A**. Examples of wild-type spermatocytes at different stages of meiotic prophase I. Heterochromatin was visualized using DAPI. SYCP3 and SYCP1 immunostaining were used to stage spermatocytes and visualize the SC at paired centromeres. CREST served as a marker for centromeres. **B**. Example of a late diplotene spermatocyte in which most centromere pairs share a common pericentromeric heterochromatin cloud. Blue arrow indicates an example of a centromere pair in a shared heterochromatin cloud. White arrowheads indicate a chromosome pair for which the pericentromeric heterochromatin is in separate clouds. Magnified chromosomes show details of pairs of homologs. **C**. Electron microscopy of silver-stained wild-type diplotene spermatocytes showing examples of chiasmate (left) and apparently achiasmate (right) chromosomes connected by electron-dense pericentromeric heterochromatin. Arrows indicate centromeres homologous pairs. **D**. Quantitation of heterochromatin and centromere association in mouse spermatocytes. Individual chromosomes in chromosome spreads from diplotene mouse spermatocytes were scored for whether their centromeres (CREST) were in the same or different heterochromatin clouds. Chiasmate pairs n=233, achiasmate pairs n=26. Statistical comparison was with Fishers exact test. Scale bars = 5 µm except for magnified images of individual chromosomes.

This association of the pericentromeric heterochromatin of diplotene centromeres was confirmed by electron microscopy of silver-stained diplotene spermatocytes (Fig. 1 C) which showed that the centromeres of homologous chromosomes, regardless of whether they underwent exchange, remain in close proximity - connected by electron-dense clouds of pericentromeric heterochromatin.

If the heterochromatin cloud can act as a tether between non-exchange partner chromosomes, then centromere pairs should remain in the same heterochromatin cloud regardless of whether the pair is tethered by a chiasma. To test this, we scored individual chiasmate and apparently-achiasmate partners in late diplotene chromosomes spreads, in which there was no more visible SC holding the centromeres together, for whether the homologous centromeres remained connected through a common heterochromatin cloud. We found there was no significant difference between the frequency of chiasmate or achaismate centromere pairs remaining in a shared heterochromatin cloud (87% vs 96% respectively; p=0.33) (Fig. 1 D), consistent with the model that the shared heterochromatin may be sufficient to keep partner centromeres joined, in the absence of chiasmata.

As an alternative way to visualize the localization of the homologous centromeres with heterochromatin, we stained chromosome spreads with antibodies against methylated histone 3 lysine 9 (H3K9Me3), a post-translational modification of constitutive pericentromeric heterochromatin (Peters et al. 2001; Lehnertz et al. 2003) (Fig. 2 A). In late diplotene chromosome spreads, homologous centromeres (identified by their SYCP3 knobs) although well-separated, were often in a cloud of shared H3K9me3-modified heterochromatin, and multiple homologous centromere pairs sometimes shared a common heterochromatin cloud, as shown previously (Berríos et al. 2010; Hopkins et al. 2014). There is considerable variation in the amount of heterochromatin at different centromeres by both DAPI and H3K9me3 staining (Fig. 2 A). If associations between blocks of peri-centromeric heterochromatin help to keep homologous centromeres together, we reasoned that those with more abundant heterochromatin might remain together more efficiently. To test this, we categorized centromeres as having abundant or weak H3K9me3 staining then measured the distances between the centromeric SYCP3 knobs (Fig. 2 A and B). Centromere pairs that were farther than 0.8 µm apart were scored as “separated” (Fig. 2 C). By this criterion, centromere pairs with low levels of heterochromatin were more likely to become separated in chromosome spreads (48% vs 27%; p <0.01), consistent with the model that associations of the homologous pericentromeric regions keeps the centromeres together.

**Figure 2.**
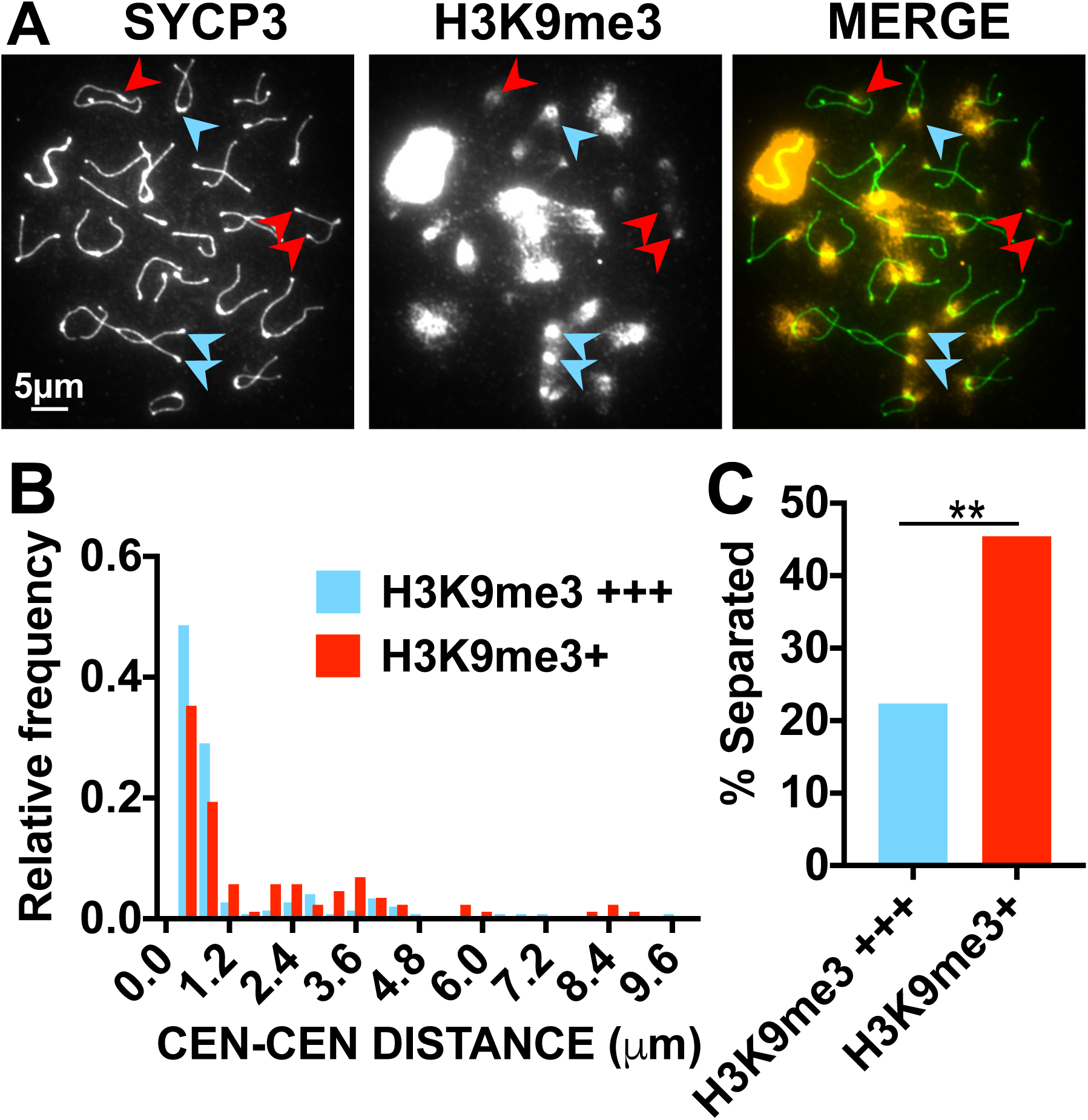
Centromeres with more heterochromatin are more likely to remain together. **A**. Chromosome spreads were stained with antibodies against H3K9me3 to mark pericentromeric heterochromatin and SYCP3 to identify chromosome axes. Chromosomes were categorized as having bright or dim H3K9me3 staining and the distances between the SYCP3 centromere knobs was measured. Blue and red arrowheads indicate examples bright and dim H3K9me3 staining. **B**. Distances between centromere pairs (in 0.4 µm bins). n=144 bright centromere pairs, 88 dim centromere pairs. **C**. Centromeres farther apart than 0.8 µm were scored as separated. Centromere pairs with dim (+) H3K9me3 staining were significantly more likely to be separated than those with bright (+++) H3K9me3 staining (Fishers exact test, p=0.0016).

### Homologous Peri-centromeric Associations are Formed in late Prophase

Previous studies have suggested that centromeres move from heterologous pericentromeric clusters to homologous centromere pairing in zygotene (Takada et al. 2011). To confirm this, we tracked the behavior of the centromeres from chromosomes 2, 8, and 15 in cells at different stages of meiotic prophase (Fig. S2). We found that in early prophase (leptotene) homologous centromeres were nearly always in different chromocenters, but by pachytene they were tightly paired in the same chromocenter. Thus, the clustering of centromeres in the chromocenters appears not to be based on homology. The clustering of peri-centric heterochromatic regions is reminiscent of the homology-independent “centromere coupling” phenomena that occurs in early meiotic prophase in several organisms ((Tsubouchi 2005), reviewed in (Obeso et al. 2014).

These results suggest the model that synapsis drives the re-organization of the pericentromeric heterochromatin into homologous clusters. Consistent with this model, previous studies of chromosome synapsis in mouse spermatocytes revealed that centromere regions are often the last to synapse (Bisig et al. 2012). To test this, we evaluated whether partially synapsed homologous partners in zygotene cells (cells that are undergoing chromosome synapsis) have their centromeres in different or the same chromocenter (Fig. 3 A and B). Individual chromosomes were scored for the ratio of the length of synaptonemal complex (SYCP1) versus the length of the chromosome axis (SYCP3). Chromosomes in the very early stages of synapsis (with SC extending only 2/5 the length of the axis) the homologous centromeres were rarely in the same chromocenter (Fig. 3 B) and as the SC increased to full-length (in pachytene) the centromeres moved completely into shared chromocenters (Fig. 3 A and B). Diplotene cells, in which SC was completely disassembled, continued to show homologous centromeres sharing chromocenters, though sometimes they shared the chromocenters with another homologous pair (Fig. 3 A and B).

**Figure 3.**
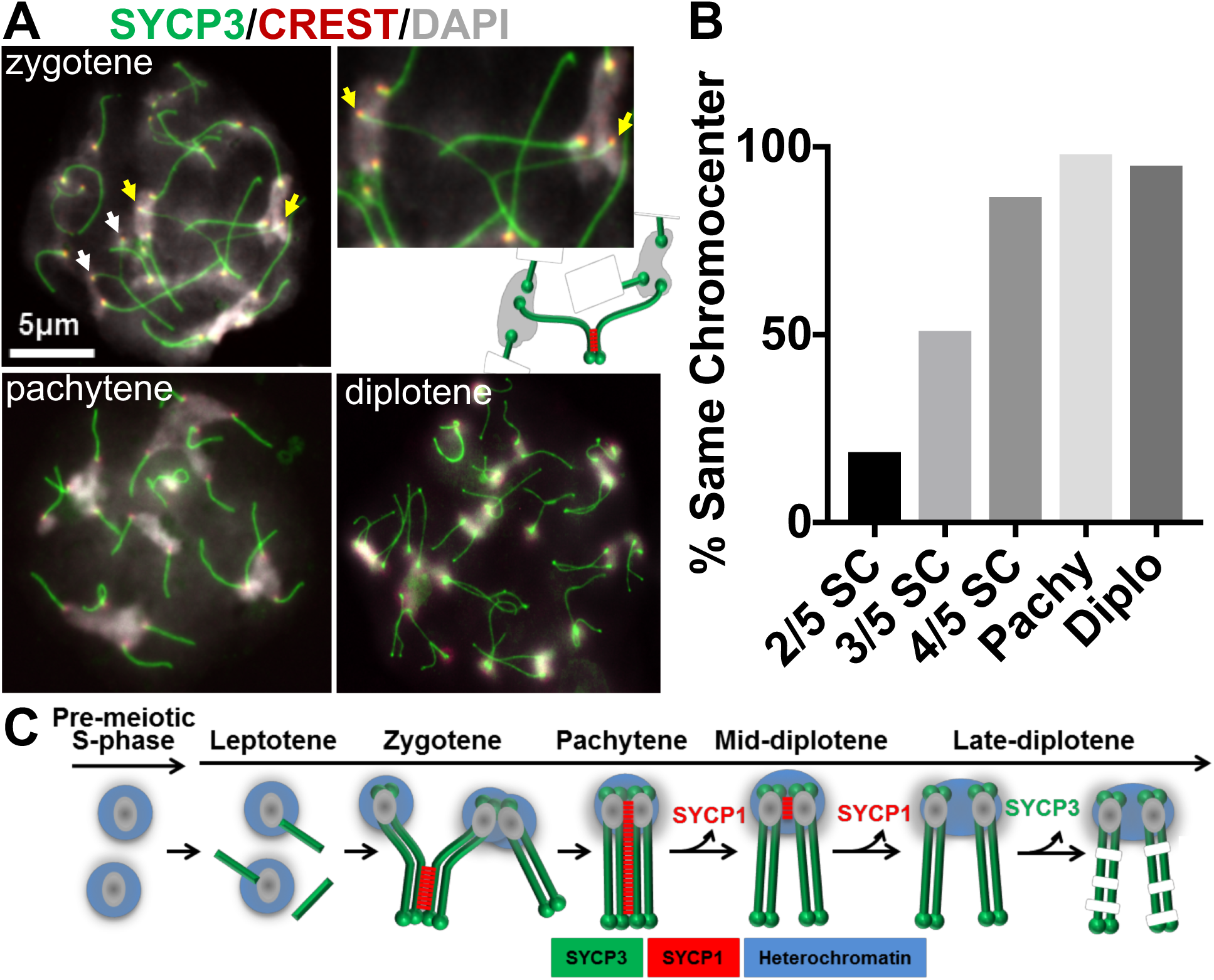
Pericentric associations move from non-homologous to homologous in meiotic prophase. **A**. Representative chromosome spreads from spermatocytes in different stages of meiotic prophase. Staging was deterimined by the amount of synapsis. Spreads were stained to visualize chromosome axes (SYCP3; green), centromeres (CREST; red) and chromatin (DAPI; gray). In zygotene spreads unsynapsed centromeres are frequently in different chromocenters. Yellow and white arrows indicate two examples of centromere pairs in different chromocenters. The magnified inset shows a centromere pair (yellow arrows) in different chromocenters. The cartoon illustrates the organization of this centromere pair. In the cartoon, green represents chromosome axes and red represents SC which is not stained in the chromosome spreads. In pachytene homologous centromere pairs are usually in the same chromocenters but often with other centromere pairs. In diplotene spreads Chromocenters are often smaller with only one or two centromere pairs.. Scale bar = 5 µm for all images except magnified images of individual chromosomes. **B**. The individual chromosome pairs in zygotene, pachytene and diplotene spreads were scored for whether the homologous centromeres were in the same chromocenters. n=100 chromosomes for each meiotic stage. **C**. Cartoon summarizing the behavior of homologous pericentromeric regions during meiotic prophase.

### Role for the SC in Establishing Homologous Pericentromeric Heterochromatin Connections

The above results suggest the model (Fig. 3 C) that SC assembly helps to pull homologous centromeres out of different chromocenters and allows the formation of smaller chromocenters harboring one or two homologous centromere pairs.

This model predicts that in mutants defective in synapsis, the non-homologous centromeric clustering will not be driven into homologous centromere associations. To test this, we analyzed the dependence of homologous peri-centromeric associations on SYCP1 (Fig. 4 A). Chromosome spreads from wild-type and *Sycp1*^-/-^ spermatocytes that exhibited diplotene-like chromosome morphologies, were scored for whether homologous centromere pairs were in the same or different chromocenters. In wild-type cells the vast majority of homologous chromosomes shared a common chromocenter (Fig. 4 A and B), only 14 of 500 chromosomes scored (3%) had their homologous centromeres separated into different chromocenters. In the *Sycp1*^-/-^ spermatocytes, the homologous chromosome axes become aligned (Fig. 4 A), as described previously (de Vries et al., 2005). However, in sharp contrast to wild-type cells we observed a high number of *Sycp1^-/-^* spermatocytes in which the centromeres of the two homologs were in different chromocenters (135 of 290 (47%) chromosomes scored) (Fig.4 A and B). We conclude that in *Sycp1*^-/-^ mutants, pericentromeric regions do not undergo the heterologous to homologous transition, even though the chromosome axes do become aligned.

**Figure 4.**
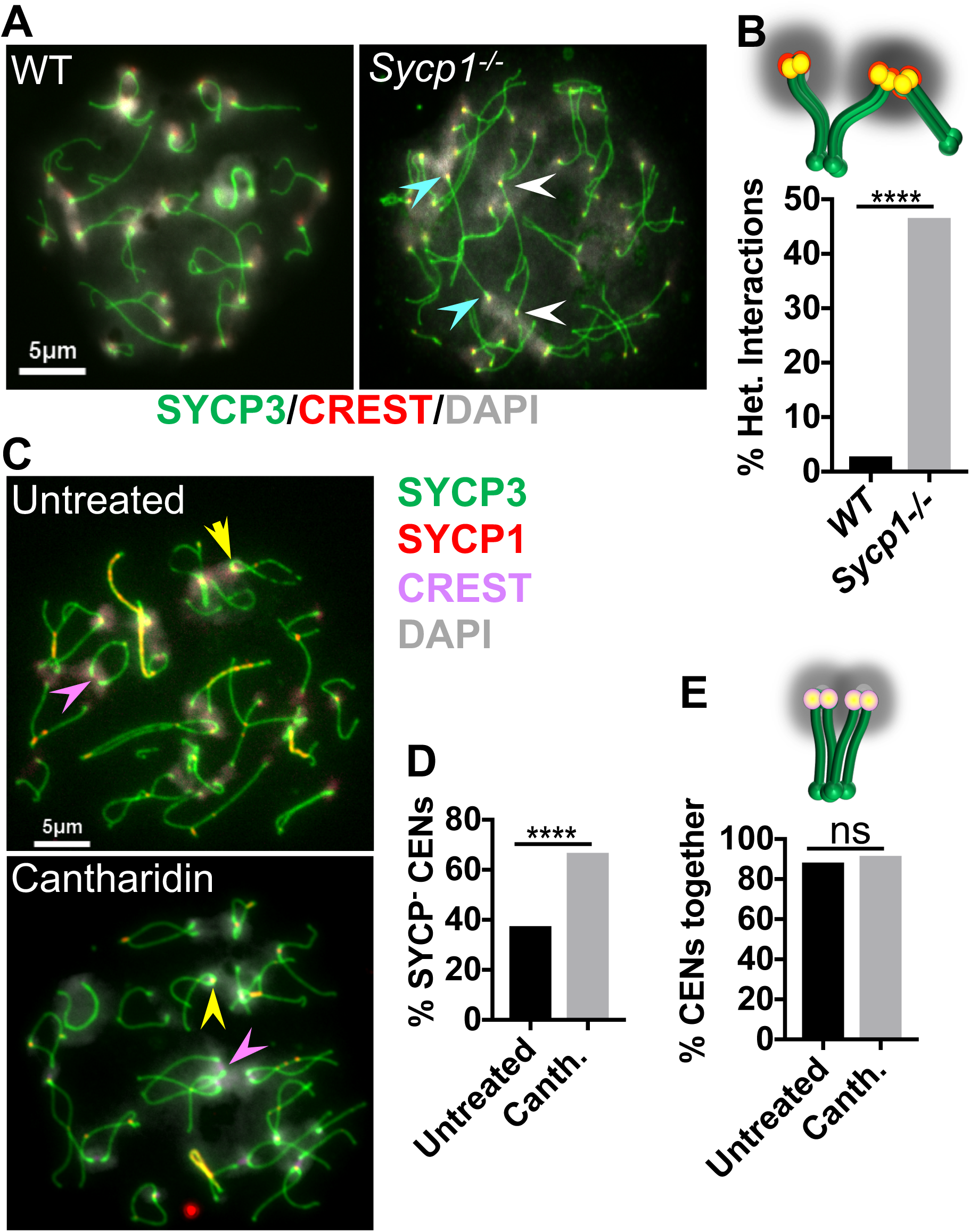
The SC is required for establishment but not maintenance of homologous heterochromatin-mediated centromere connections. **A**. Chromosome spreads from diplotene wild-type and *Sycp1^-/-^* dpermatocytes were evaluated to evaluate the role of synapsis in the merging of homologous pericentromeric regions into a shared chromocenter. Cells were stained with DAPI (grey) and antibodies against SYCP3 (green) and CREST (red). Blue and white arrowheads indicate examples of two homologous centromere pairs that are in separate chromocenters. **B**. Quantification of the frequency of chromosomes that had their centromeres in chromocenters with heterologous partners rather than in the same chromocenter. n= 500 chromosomes for wild-type and 290 for *Sycp1^-/-^*. p<0.0001, Fishers Exact test. **C**. Examples of chromosome spreads from cultured diplotene spermatocytes, with or without a three hour treatment with 30 µM cantharidin. Yellow arrowheads indicate examples of cells with persisting SYCP1 mediating centromere pairing. Pink arrowheads indicate examples of homologous centromere pairs with no detectable SYCP1. **D**. Quantification of the percentage of chromosome pairs that were negative for SYCP1 (SYCP1^-^). n=160 chromosomes from both untreated and cantharidin treated spermatocytes. p<0.0001, Fishers Exact test. **E**. Quantification of the percent of homologous centromere pairs that were together (sharing the same chromocenter) in chromosome spreads from untreated or cantharidin treated spermatocytes. n=107 centromere pairs for untreated and 60 centromere pairs for cantharidin treated. p=0.59, Fishers Exact test. Scale bars = 5 µm.

Once the homologous pericentromeric regions become aligned through synapsis, is SYCP1 still required for the persistence of the pericentromeric associations? To test this, we took advantage of previous studies showing that inhibition of PP2A phosphatase drives SC disassembly in cultured spermatocytes (Wiltshire et al. 1995). SC disassembly is driven in part by phosphorylation (Tarsounas et al. 1999; Sourirajan and Lichten 2008; Sun and Handel 2008; Jordan et al. 2012; Argunhan et al. 2017), reviewed in (Gao and Colaiácovo 2018) and PP2A presumably acts in prophase to reverse critical phosphorylations that drive SC disassembly. We treated cultured spermatocytes with the phosphatase inhibitor cantharidin (Honkanen 1993), then examined chromosome spreads to determine first, if cantharidin promotes the loss of the persistent SC at paired centromeres in diplotene spermatocytes (Fig. 4 C). In diplotene cells, cantharidin treatment significantly increased the numbers of centromere pairs without SYCP1 (Fig. 4 D; SYCP1^-^ CENs). Thus, cantharidin treatment in diplotene is causes the loss of centromeric SYCP1 after the centromeres have been paired. This allowed us to ask whether centromeres that are no longer directly tethered by SC continued to share a common chromocenter. The loss of SYCP1 did not result in a separation of the centromere pairs, instead, they remained joined by a shared cloud of heterochromatin (Fig. 4 E). These results indicate that heterochromatin connections have been already established by mid-late diplotene, when SYCP1 remains at the centromere to mediate centromere pairing. The results also indicate that heterochromatin connections between homolog centromeres are stable in the absence of a centromeric SC, raising the possibility that these connections might provide a link between homologous centromeres that could contribute to their bi-orientation as they transition from prophase into pro-metaphase.

## Discussion

### Role for Heterochromatin in Maintaining Meiotic Chromosome Alignment

In *Drosophila* females pericentromeric chromatin has been implicated in helping promote the segregation of homologous chromosomes, even if they fail to be joined by chiasmata (Karpen et al. 1996; Dernburg et al. 1996). In Drosophila females, chromosomes that fail to undergo recombination (*X* and *4*) are connected by heterochromatic threads during prometaphase I as chromosomes orient on the meiotic spindle. These threads have been proposed to serve as a connection between the partners that may help them to bi-orient on the spindle (Hughes et al. 2009). Additional evidence for the conservation of heterochromatic threads connecting chromosomes during meiosis comes from *Drosophila* and crane fly sperm (LaFountain et al. 2002; Hartl et al. 2008). The results presented here demonstrate that heterochromatin also plays a role in promoting meiotic centromere interactions in the mouse, and that these interactions are consistent with a role in promoting proper meiotic segregation, especially of achiasmate partners.

### Origin and Regulation of Heterochromatin-Mediated Centromere Clustering Early in Prophase

Observations in a wide range of organisms show that very early in the meiotic program (leptotene), before homologous pairing occurs, centromeres associate in pairs or clusters independent of sequence homology (reviewed in (Obeso et al. 2014). This is termed centromere coupling (Tsubouchi 2005). This work confirms previous observations of centromere clustering in the mouse suggesting that it resembles centromere coupling (Berríos et al. 2010). Our results show that as cells progress through prophase, centromeres move from large chromocenters bearing multiple heterologous centromeres to smaller chromocenters that include their homologous partners. The mechanism by which the pericentric heterochromatic regions become re-organized has not been clear, but our results suggest that it is driven by synapsis. First, the centromeres move begin sharing their chromocenters with their homologous partners as the synaptonemal complex lengthens. Second, in mutants that are incapable of synapsis, the homologous axes still align, but the centromeres remain in chromocenters with non-homologous partners. However, synapsis cannot be the only mechanism controlling this chromocenter re-organization. After complete synapsis in pachytene, in the transition to diplotene, the chromocenters continue to individualize, moving from clumps of homologous centromere pairs to mostly single homologous centromere pairs. It is not known what drives this resolution of the centromere clusters to individual pairs. But the pericentric heterochromatin in the chromocenters is rich in cohesin, condensin, and topoisomerase II (Ishiguro et al. 2011; Verver et al. 2013; Gómez et al. 2013; Ishiguro et al. 2014; Hopkins et al. 2014). It may be that the interplay of these chromatin compaction factors is important for regulating the formation and dissolution of chromocenters. Consistent with this notion, mutation of the cohesin gene *Stag3* in mice increases the number of chromocenters suggesting that cohesins are necessary for holding together the pericentric heterochromatin of multiple chromosomes (Hopkins et al. 2014).

### Origin and Regulation of Homologous Heterochromatin Connections

When and how are homologous heterochromatin connections established? Our results define a period of prophase I in which the SC promotes the stable homologous pericentromeric heterochromatin interactions observed between diplotene chromosomes. First, prior to synapsis spermatocytes display high numbers of non-homologous centromeres connected by heterochromatin. Second, diplotene *Sycp1^-/-^* spermatocytes have abnormally high numbers of unpaired chromosomes and chromosomes engaged in non-homologous centromeric associations, suggesting the SC plays a role in establishment of homologous heterochromatin connections. However, once homologous centromeres have been juxtaposed by synapsis, the SC is no longer necessary to maintain the association of the homologous pericentromeric heterochromatin regions, since precocious removal of the SC from paired centromeres of diplotene chromosomes (using a PP2A inhibitor) disrupted the close juxtaposition of homologous centromeres (centromere pairing) but did not affect heterochromatin interactions between homologous pairs. Thus, there must be a mechanism that stabilizes heterochromatin connections between homolog pairs independently of the SC.

What is the nature of heterochromatin interactions and what activity could disrupt heterochromatin connections from homologous centromeres when they segregate? The tight physical association of heterochromatin observed in Drosophila oocytes during early meiosis suggested the possibility that heterochromatin connections may be established during DNA replication (Dernburg et al. 1996). It has been suggested that linkages are established during stalled replication fork repair (Hughes et al. 2009), however our results suggest that the coalescence of pericentric heterochromatin into chromocenters containing multiple centromere does not happen until well after S-phase.

The persistence of heterochromatic associations into meiotic pro-metaphase is reminiscent of the ultra-fine DNA threads that connect sister chromatids in mitotic cells (Chan et al. 2009). The connections between mitotic sister chromatid DNAs that are responsible for these threads occur through multiple mechanisms including catenation, late replication intermediates, and telomere fusion events (Liu et al. 2014).

It is possible that protein-protein or protein-DNA interactions of a different nature may promote post-pachytene stable homologous heterochromatin interactions. Centromeric regions are enriched for cohesion proteins and the roles of different types of meiotic cohesion complexes remain unclear. It is possible that cohesins act to form interhomolog cohesion that links centromeric heterochromatin, or alternatively, provide an environment in which catenation or other links between partner chromosomes are maintained until metaphase. Such a mechanism for linking homologous heterochromatic regions would require a novel meiotic remodeling of cohesins as cells move through prophase. This would include dissolving cohesive links between non-homologous heterochromatin blocks and establishing cohesion between homologous heterochromatin blocks after they are brought together by SC formation. Further work will be necessary to test these hypotheses.

## Experimental Procedures

### Mouse Strains

The Oklahoma Medical Research Foundation Animal Care and Use Committee (IACUC) approved all animal protocols. Wild type (C57BL/6) and *Sycp1^-/-^* mice (de Vries *et al*., 2005) were used in this study.

### Cytology

We employed established experimental approaches for the visualization of chromosomes in chromosome surface spreads (Peters et al. 1997). Incubations with primary antibodies were carried out for 12 h at 4°C in 1× PBS plus BSA 2%. To detect SYCP1 and SYCP3 we used polyclonal antibodies raised against mouse SYCP1 at 1:150 dilution (Novus Biologicals, NB300–229) and polyclonal chicken antibody generated in our laboratory raised against mouse SYCP3, at 1:300 dilution. Centromeres were detected using the human centromere protein antibody (CREST, Antibody Incorporated, 9101–02) at 1:50 dilution. H3K9me3 was detected using polyclonal rabbit antibody raised against H3K9me3 at 1:500 dilution. Following three washes in 1× PBS, slides were incubated for 1 hour at room temperature with secondary antibodies. A combination of Fluorescein isothiocyanate (FITC)-conjugated goat anti-rabbit IgG (Jackson laboratories) with Rhodamine-conjugated goat anti-mouse IgG and Cy5-conjugated goat anti-human IgG each diluted 1:350 were used for simultaneous immunolabeling if required. Slides were subsequently counterstained for 3 min with 2 µg/ml DAPI containing Vectashield mounting solution (Vector Laboratories) and sealed with nail varnish. We used an Axiovision SE 64 (Carl Zeiss, inc.) for imaging acquisition and processing.

Spermatocyte chromosome spreads for electron microscopy analysis was performed as previously described (Dresser et al. 1987).

### Spermatocyte culturing and chemical inhibition

Short-term culture of spermatocytes was performed essentially as described (La Salle et al. 2009). Cantharidin was added at 30 µM (Millipore; 505156; 30 mM stock dissolved in DMSO) and incubated for 3 hours. Cells were then pelleted, washed with 1X PBS and processed for surface spreads. Equivalent volumes of DMSO were added to “no treatment” control cultures.

### FISH combined with immunostaining

DNA FISH was carried out essentially as previously described (Turner et al. 2005). Cell suspensions were prepared in 1X PBS containing a cocktail of protease inhibitors. Cells were spun down and resuspended in 100 mM sucrose, pH 7.2. Approximately 70 μl of this cell suspension was dropped on clean slides and allowed to attach to the slides for 10 min at RT. Slides were immersed in 4% paraformaldehyde for 20 min and rinsed in 1X PBS, dehydrated through an ethanol series (2 × 70%, 80%, 96%, 100%) and air-dried. Hybridization solution specific fluorescent point probes for chromosomes 2, 8, and 15 were obtained from ID Labs Inc. Samples were incubated in humid chambers for 24h at 37 ºC. We then subjected slides to washes at 42 ºC (three washes with 2x SSC and 50% formamide and three washes with 2x SSC) and transferred them to 4xSSC and 0.1% Tween-20. Slides were blocked in 4x SSC, 4 mg/ml bovine serum albumin and 0.001% Tween-20, for 30 min at 37 ºC. At each of these steps, the slides were incubated for 30 min at 37 ºC and washed three times for 2 min each in 4xSSC and 0.1% Tween-20. Slides were cross-linked with 1%PFA/1xPBS for 10 min and immunostained with the corresponding antibody.

### Statistical tests

The statistical tests were used are described in the text and Figure legends. Statistical tests were performed using Prizm software.

## Supporting information

## Supplemental Information

Supplemental Information includes two figures and can be found with this article online.

## Acknowledgements

This work was supported by grant R01 GM125803 to RJP, and NIH grant R01 GM087377 to DSD. Electron microscopy was performed in the OMRF Imaging Core with their technical support. Sycp1^-/-^ mice were a gift from Christer Höög.

## References

Argunhan B, Leung W-K, Afshar N, et al (2017) Fundamental cell cycle kinases collaborate to ensure timely destruction of the synaptonemal complex during meiosis. The EMBO Journal 36:2488–2509. doi: 10.15252/embj.201695895

Bascom-Slack C, Ross LO, Dawson DS (1997) 7. Chiasmata, Crossovers, and Meiotic Chromosome Segregation. Advances in Genetics 35:253–284. doi: 10.1016/S0065-2660(08)60452-6

Baudat F, Manova K, Yuen JP, et al (2000) Chromosome synapsis defects and sexually dimorphic meiotic progression in mice lacking Spo11. Mol Cell 6: 989–998.

Berríos S, Manieu C, López-Fenner J, et al (2014) Robertsonian chromosomes and the nuclear architecture of mouse meiotic prophase spermatocytes. Biol Res 47:16. doi: 10.1186/0717-6287-47-16

Berríos S, Manterola M, Prieto Z, et al (2010) Model of chromosome associations in Mus domesticus spermatocytes. Biol Res 43: 275–285.

Bisig CG, Guiraldelli MF, Kouznetsova A, et al (2012) Synaptonemal complex components persist at centromeres and are required for homologous centromere pairing in mouse spermatocytes. PLoS Genet 8:e1002701. doi: 10.1371/journal.pgen.1002701

Chan K-L, Palmai-Pallag T, Ying S, Hickson ID (2009) Replication stress induces sister-chromatid bridging at fragile site loci in mitosis. Nat Cell Biol 11:753–760. doi: 10.1038/ncb1882

Cheng EY, Hunt PA, Naluai-Cecchini TA, et al (2009) Meiotic recombination in human oocytes. PLoS Genet 5:e1000661. doi: 10.1371/journal.pgen.1000661

Davis L, Smith GR (2003) Nonrandom homolog segregation at meiosis I in Schizosaccharomyces pombe mutants lacking recombination. Genetics 163: 857–874.

Dawson DS, Murray AW, Szostak JW (1986) An alternative pathway for meiotic chromosome segregation in yeast. Science 234: 713–717.

Dernburg AF, McDonald K, Moulder G, et al (1998) Meiotic recombination in C. elegans initiates by a conserved mechanism and is dispensable for homologous chromosome synapsis. Cell 94: 387–398.

Dernburg AF, Sedat JW, Hawley RS (1996) Direct evidence of a role for heterochromatin in meiotic chromosome segregation. Cell 86: 135–146.

Ding D-Q, Yamamoto A, Haraguchi T, Hiraoka Y (2004) Dynamics of homologous chromosome pairing during meiotic prophase in fission yeast. Dev Cell 6: 329–341.

Dresser M, Pisetsky D, Warren R, et al (1987) A new method for the cytological analysis of autoantibody specificities using whole-mount, surface-spread meiotic nuclei. J Immunol Methods 104: 111–121.

Fledel-Alon A, Wilson DJ, Broman K, et al (2009) Broad-scale recombination patterns underlying proper disjunction in humans. PLoS Genet 5:e1000658. doi: 10.1371/journal.pgen.1000658

Gao J, Colaiácovo MP (2018) Zipping and Unzipping: Protein Modifications Regulating Synaptonemal Complex Dynamics. Trends Genet 34:232–245. doi: 10.1016/j.tig.2017.12.001

Gladstone MN, Obeso D, Chuong H, Dawson DS (2009) The synaptonemal complex protein Zip1 promotes bi-orientation of centromeres at meiosis I. PLoS Genet 5:e1000771. doi: 10.1371/journal.pgen.1000771

Gómez R, Jordan PW, Viera A, et al (2013) Dynamic localization of SMC5/6 complex proteins during mammalian meiosis and mitosis suggests functions in distinct chromosome processes. J Cell Sci 126:4239–4252. doi: 10.1242/jcs.130195

Grell RF (1962) A new model for secondary nondisjunction: the role of distributive pairing. Genetics 47: 1737–1754.

Hartl TA, Sweeney SJ, Knepler PJ, Bosco G (2008) Condensin II resolves chromosomal associations to enable anaphase I segregation in Drosophila male meiosis. PLoS Genet 4:e1000228. doi: 10.1371/journal.pgen.1000228

Hawley RS, Irick H, Zitron AE, et al (1992) There are two mechanisms of achiasmate segregation in Drosophila females, one of which requires heterochromatic homology. Dev Genet 13:440–467. doi: 10.1002/dvg.1020130608

Honkanen RE (1993) Cantharidin, another natural toxin that inhibits the activity of serine/threonine protein phosphatases types 1 and 2A. FEBS Letters 330: 283–286.

Hopkins J, Hwang G, Jacob J, et al (2014) Meiosis-specific cohesin component, Stag3 is essential for maintaining centromere chromatid cohesion, and required for DNA repair and synapsis between homologous chromosomes. PLoS Genet 10:e1004413. doi: 10.1371/journal.pgen.1004413

Hughes SE, Gilliland WD, Cotitta JL, et al (2009) Heterochromatic Threads Connect Oscillating Chromosomes during Prometaphase I in Drosophila Oocytes. 5:e1000348. doi: 10.1371/journal.pgen.1000348.s016

Ishiguro K-I, Kim J, Fujiyama-Nakamura S, et al (2011) scientific report. EMBO reports 12:1–9. doi: 10.1038/embor.2011.2

Ishiguro K-I, Kim J, Shibuya H, et al (2014) Meiosis-specific cohesin mediates homolog recognition in mouse spermatocytes. Genes Dev 28:594–607. doi: 10.1101/gad.237313.113

Jordan PW, Karppinen J, Handel MA (2012) Polo-like kinase is required for synaptonemal complex disassembly and phosphorylation in mouse spermatocytes. J Cell Sci 125:5061–5072. doi: 10.1242/jcs.105015

Karpen GH, Le MH, Le H (1996) Centric heterochromatin and the efficiency of achiasmate disjunction in Drosophila female meiosis. Science 273: 118–122.

Kemp B, Boumil RM, Stewart MN, Dawson DS (2004) A role for centromere pairing in meiotic chromosome segregation. Genes Dev 18:1946–1951. doi: 10.1101/gad.1227304

Klapholz S, Waddell CS, Esposito RE (1985) The role of the SPO11 gene in meiotic recombination in yeast. Genetics 110: 187–216.

Klein F, Mahr P, Galova M, et al (1999) A central role for cohesins in sister chromatid cohesion, formation of axial elements, and recombination during yeast meiosis. Cell 98:91–103. doi: 10.1016/S0092-8674(00)80609-1

La Salle S, Sun F, Handel MA (2009) Isolation and short-term culture of mouse spermatocytes for analysis of meiosis. Methods Mol Biol 558:279–297. doi: 10.1007/978-1-60761-103-5_17

LaFountain JR, Cole RW, Rieder CL (2002) Partner telomeres during anaphase in crane-fly spermatocytes are connected by an elastic tether that exerts a backward force and resists poleward motion. J Cell Sci 115: 1541–1549.

Lehnertz B, Ueda Y, Derijck AAHA, et al (2003) Suv39h-mediated histone H3 lysine 9 methylation directs DNA methylation to major satellite repeats at pericentric heterochromatin. Curr Biol 13: 1192–1200.

Liu Y, Nielsen CF, Yao Q, Hickson ID (2014) The origins and processing of ultra fine anaphase DNA bridges. Current Opinion in Genetics & Development 26:1–5. doi: 10.1016/j.gde.2014.03.003

Martens JHA, O’Sullivan RJ, Braunschweig U, et al (2005) The profile of repeat-associated histone lysine methylation states in the mouse epigenome. EMBO J 24:800–812. doi: 10.1038/sj.emboj.7600545

Moens PB, Spyropoulos B (1995) Immunocytology of chiasmata and chromosomal disjunction at mouse meiosis. Chromosoma 104: 175–182.

Mouse Genome Sequencing Consortium, Waterston RH, Lindblad-Toh K, et al (2002) Initial sequencing and comparative analysis of the mouse genome. Nature 420:520–562. doi: 10.1038/nature01262

Newnham L, Jordan P, Rockmill B, et al (2010) The synaptonemal complex protein, Zip1, promotes the segregation of nonexchange chromosomes at meiosis I. Proceedings of the National Academy of Sciences 107:781–785. doi: 10.1073/pnas.0913435107

Nicklas RB (1997) How Cells Get the Right Chromosomes. 275:632–637. doi: 10.1126/science.275.5300.632

Nicklas RB, Koch CA (1969) Chromosome micromanipulation. 3. Spindle fiber tension and the reorientation of mal-oriented chromosomes. J Cell Biol 43: 40–50.

Obeso D, Pezza RJ, Dawson D (2014) Couples, pairs, and clusters: mechanisms and implications of centromere associations in meiosis. Chromosoma 123:43–55. doi: 10.1007/s00412-013-0439-4

Oliver TR, Feingold E, Yu K, et al (2008) New Insights into Human Nondisjunction of Chromosome 21 in Oocytes. 4:e1000033. doi: 10.1371/journal.pgen.1000033.s001

Pardue ML, Gall JG (1970) Chromosomal localization of mouse satellite DNA. Science 168: 1356–1358.

Parra MT (2004) Involvement of the cohesin Rad21 and SCP3 in monopolar attachment of sister kinetochores during mouse meiosis I. J Cell Sci 117:1221–1234. doi: 10.1242/jcs.00947

Peters AH, O’Carroll D, Scherthan H, et al (2001) Loss of the Suv39h histone methyltransferases impairs mammalian heterochromatin and genome stability. Cell 107: 323–337.

Peters AH, Plug AW, van Vugt MJ, de Boer P (1997) A drying-down technique for the spreading of mammalian meiocytes from the male and female germline. Chromosome Res 5: 66–68.

Qiao H, Chen JK, Reynolds A, et al (2012) Interplay between Synaptonemal Complex, Homologous Recombination, and Centromeres during Mammalian Meiosis. 8:e1002790. doi: 10.1371/journal.pgen.1002790.s002

Romanienko PJ, Camerini-Otero RD (2000) The mouse Spo11 gene is required for meiotic chromosome synapsis. Mol Cell 6: 975–987.

Sansam CL, Pezza RJ (2015) Connecting by breaking and repairing: mechanisms of DNA strand exchange in meiotic recombination. FEBS J 282:2444–2457. doi: 10.1111/febs.13317

Scherthan H, Weich S, Schwegler H, et al (1996) Centromere and telomere movements during early meiotic prophase of mouse and man are associated with the onset of chromosome pairing. J Cell Biol 134: 1109–1125.

Schwarzacher T, Mayr B, Schweizer D (1984) Heterochromatin and nucleolus-organizer-region behaviour at male pachytene of Sus scrofa domestica. Chromosoma 91: 12–19.

Sourirajan A, Lichten M (2008) Polo-like kinase Cdc5 drives exit from pachytene during budding yeast meiosis. Genes Dev 22:2627–2632. doi: 10.1101/gad.1711408

Sun F, Handel MA (2008) Regulation of the meiotic prophase I to metaphase I transition in mouse spermatocytes. Chromosoma 117:471–485. doi: 10.1007/s00412-008-0167-3

Takada Y, Naruse C, Costa Y, et al (2011) HP1γ links histone methylation marks to meiotic synapsis in mice. Development 138:4207–4217. doi: 10.1242/dev.064444

Takeo S, Lake CM, Morais-de-Sá E, et al (2011) Synaptonemal Complex-Dependent Centromeric Clustering and the Initiation of Synapsis in Drosophila Oocytes. Current Biology 21:1845–1851. doi: 10.1016/j.cub.2011.09.044

Tarsounas M, Pearlman RE, Moens PB (1999) Meiotic activation of rat pachytene spermatocytes with okadaic acid: the behaviour of synaptonemal complex components SYN1/SCP1 and COR1/SCP3. J Cell Sci 112 (Pt 4):423–434.

Tease C, Hartshorne GM, Hultén MA (2002) Patterns of meiotic recombination in human fetal oocytes. Am J Hum Genet 70:1469–1479. doi: 10.1086/340734

Tsubouchi T (2005) A Synaptonemal Complex Protein Promotes Homology-Independent Centromere Coupling. 308:870–873. doi: 10.1126/science.1108283

Turner JMA, Mahadevaiah SK, Fernandez-Capetillo O, et al (2005) Silencing of unsynapsed meiotic chromosomes in the mouse. Nat Genet 37:41–47. doi: 10.1038/ng1484

Verver DE, van Pelt AMM, Repping S, Hamer G (2013) Role for rodent Smc6 in pericentromeric heterochromatin domains during spermatogonial differentiation and meiosis. Cell Death Dis 4:e749. doi: 10.1038/cddis.2013.269

Wiltshire T, Park C, Caldwell KA, Handel MA (1995) Induced premature G2/M-phase transition in pachytene spermatocytes includes events unique to meiosis. Developmental Biology 169:557–567. doi: 10.1006/dbio.1995.1169

Woods LM, Hodges CA, Baart E, et al (1999) Chromosomal influence on meiotic spindle assembly: abnormal meiosis I in female Mlh1 mutant mice. J Cell Biol 145: 1395–1406.

