## Supplementary material for "Heterochromatin Interactions Maintain Homologous Centromere Associations in Mouse Spermatocyte Meiosis"

**Figure S1. Pattern of pericentromeric organization through meiotic prophase.** Squash preparations like those shown in Figure 1 A were scored to determine the timing of pericentromeric chromatin re-organization events in early meiosis. Cells were staged according to their nuclear and chromosomal morphologies. **A.** The numbers of chromocenters per nucleus were scored. Chromocenters are large and in small numbers in leptotene in significantly decrease in number as cells progress. B. The number of CREST staining foci per nucleus. As synapsis proceeds (leptotene to pachytene) the numbers of CREST foci drop significantly consistent with homologous centromere pairs being brought into close juxtaposition such that the pair yields a single CREST focus. For the above graphs, the number of nuclei scored for each timepoint are: Spermatogonia B, 53: S-phase, 72: Leptotene, 23: Zygotene, 70; Pachytene, 36; Diplotene, 47. Upaired t-tests were used for statistical comparisons. ****p<0.0001.

**Figure S2. Pre-meiotic cells and early prophase I spermatocytes show no signals for homologous pericentromeric heterochromatin connections. A**. Examples of spermatocytes showing FISH signals for a chromosome 8 region-specific probe (arrows). SYCP3 immunostaining was used for spermatocyte staging; DAPI staining was used to identify chromocenters. (**B**, **C** and **D**) Diagrams of chromosomes 8, 15 and 2 showing the FISH probe DNA target sites. Quantitation of whether the FISH foci were in the same or different chromocenters, and if the same chromocenter, whether they were paired (one signal) or unpaired (two signals). For all three loci that were evaluated, the homologous loci were normally in different chromocenters in pre-meiotic and leptotene cells, then re-organized to enter the same focus concurrent with synapsis.
